# Nectar sugar gradient within Darwin’s syndrome inflorescence shaped by yeasts rather than intrinsic plant features

**DOI:** 10.64898/2026.07.22.740096

**Authors:** Carlos M. Herrera

## Abstract

Features of floral rewards, particularly the energy content and distribution of nectar, have been traditionally considered intrinsic plant traits. Since specialized floricolous yeasts thriving in floral nectar can modify features of single flowers, they could also alter whole inflorescence traits which impinge on pollinator behavior and components of plant pollination success such as the proportion of outcrossed progeny. This study assessed yeast ability to shape variation in nectar quality within vertical inflorescences of *Gladiolus illyricus* (Iridaceae), a Darwin’s syndrome plant (flowers functionally male at first, that open and are visited by pollinators from bottom to top). In inflorescences with yeasts experimentally excluded a declining gradient in nectar sugar concentration built up from the oldest, female-stage flowers at bottom up to the youngest, male-stage ones at top. This plant-intrinsic gradient was fully reversed in inflorescences naturally exposed to yeasts. The two yeast species involved (*Metschnikowia reukaufii* and *M. gruessii*) differed in their effects on the sugar gradient. Results show that floral variation within inflorescences was shaped by flower yeast composition and abundance rather than intrinsic plant features, hence supporting the notion that some inflorescence traits typically attributed to plant genotype and pollinator-mediated selection can actually reflect the metabolism of flower microbes.

## Introduction

Ecological interactions between plants and their animal pollinators largely rely on the provision of food rewards in the form of floral nectar and pollen. The composition and energetic characteristics of such rewards, as well as their distribution in time and space, influence the foraging behavior of pollinators within and among plants, which in turn affect plant mating system and reproductive success. For example, individual pollinators probe only a fraction of the flowers available in a given plant or inflorescence depending on the mean and variance of per-flower rewards (Kadmon et al. 1991; Harder et al. 2004; Herrera 2009; Maldonado et al. 2023), and variation in the proportion of flowers probed per visit eventually translates into differences in the proportion of outcrossed progeny (Hessing 1988; Karron et al. 2004). Features of floral rewards, particularly the energy content and spatio-temporal distribution of nectar have therefore been traditionally considered as *intrinsic* plant traits subject to selection from pollinators and closely linked to the eco-evolutionary trajectories of plant-pollinator systems (Bentley and Elias 1983; Jones and Little 1983; Nicolson et al. 2007; Willmer 2011). Nevertheless, recent research has shown that floricolous microbes (yeasts and bacteria) are widespread worldwide in the nectar of entomophilous flowers exposed to pollinator visitation (Álvarez et al. 2012; Brysch-Herzberg 2004; Canto and Herrera 2012; Canto et al. 2017; de Vega et al. 2009; Herrera et al. 2008, 2009; Pozo et al. 2016), and the chemistry and rewarding value of the nectar that pollinators actually encounter while foraging in the wild do not reflect only intrinsic floral features, but also the presence, abundance, species composition and metabolism of nectar microbes. For example, the metabolism of specialized, pollinator-borne floricolous yeasts can modify nectar sugar concentration and chemical composition (Bogo et al. 2021; Canto et al. 2008; de Vega and Herrera 2013; Herrera et al. 2008; Schaeffer et al. 2015), reduce the concentration of secondary metabolites (Vanette and Fukami 2016), and alter the scent (Rering et al. 2018; Sobhy et al. 2025) and internal thermal environment (Herrera and Pozo 2010) of individual flowers, all of which can eventually modify patterns of flower visitation by pollinators (Herrera and Medrano 2017; Herrera et al. 2013; Ma et al. 2025; Schaeffer et al. 2017, 2019). These and related studies have consistently shown that yeasts thriving in floral nectar have a definite flower engineering capacity (*sensu* Jones et al. 1994), as they modify the rewards and internal environment of individual flowers in ways that can directly influence the mode in which plants interact with their pollinators.

The prevailing flower-centered perpective, which treats the individual flower as the key unit in plant-pollinator-microbe interactions, has so far produced only a partial view of the role of yeasts in plant-pollinator interactions. Whole inflorescence traits can be more influential on pollinator behavior and pollination success than single flower traits (Harder et al. 2000, 2004; Ishii et al. 2008; Morales et al. 2013). This happens, for instance, in vertical inflorescences exhibiting the so-called Darwin’s inflorescence syndrome, where flowers develop from the bottom up (acropetalous), are functionally male before female (protandrous), and are mostly pollinated by bees that typically move from the bottom to the top of the inflorescence. The predictable bottom-to-top directionality of pollinators, along with the contrasting distribution of female- (bottom) and male-stage (top) flowers in the inflorescence, will favor cross-pollination (Harder et al. 2004; McKone et al. 1995). Named after his work on *Spiranthes* orchids (Darwin 1877, pp. 113-114), the Darwin’s inflorescence syndrome has become a favored study system to elucidate the influence of whole inflorescence traits, such as gradients in nectar rewards and flower sex expression, on plant-pollinator interactions (Corbet et al. 1981; Dreisig 1989; Harder et al. 2004; Orth and Waddington 1997; Pyke 1978; Valtueña et al. 2013; Waddington and Heinrich 1979). That study system provides also an excellent opportunity for testing the engineering effects of nectar yeasts at the whole inflorescence level. This note examines the influence of nectar yeasts on the nectar quality gradient within the linear inflorescences of *Gladiolus illyricus* (Iridaceae), a long-known Darwin’s syndrome plant (Wilson 1878). Experiments reported here were inspired by previous findings showing that density of yeasts in floral nectar tended to increase, and nectar sugar concentration to decrease, with increasing flower age (Herrera et al. 2008). Results will show that the natural gradient within *G. illyricus* inflorescences in quality of nectar reward is better explained by flower yeasts than by intrinsic plant features.

## Materials and methods

*Gladiolus illyricus* is a spring-blooming plant producing vertical inflorescences with 4-8 flowers that open sequentially from bottom to top (Figure 1A). When exposed to natural pollinator visitation, individual flowers last for 2-(3) days, and inflorescences bear 2-5 flowers open simultaneously. Flowers are protandrous and produce concentrated (median = 29%, interquartile range = 23–41% sucrose equivalents, data from the present study), sucrose-dominated (Herrera 2014) nectar. Main pollinators in the study area (see below) are large bees (*Amegilla*, *Anthophora*, *Bombus*, *Eucera*, *Xylocopa*), accounting for 58% of flower visits. As noted by Wilson (1878, p. 509) and confirmed by my own observations during pollinator censuses (Herrera 2026), bees consistently begin foraging by the lowermost flower in each inflorescence.

**Figure 1.**
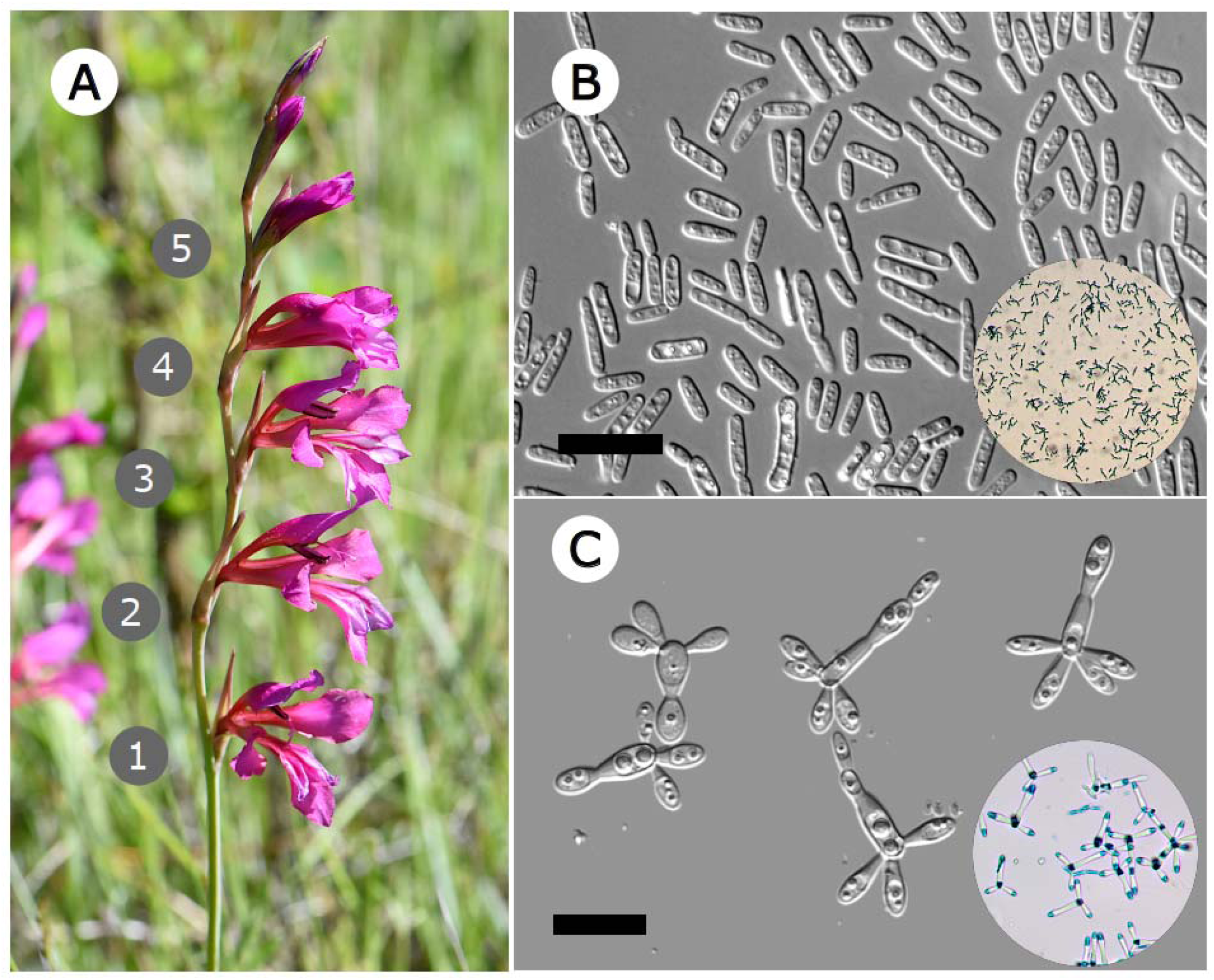
Inflorescence of *Gladiolus illyricus* with indication of nodal positions of flowers (A), and Nomarsky interference contrast photomicrographs of living *Metschnikowia reukaufii* (B) and *Metschnikowia gruessii* (C) yeast cells in floral nectar (scale bar = 25 µm). Insets show microscopic appearance of the two yeast species after fixation and staining with lactophenol cotton blue. Note the distinct pattern of staining of cells of the two species mentioned in text.

This study was carried out between 20-31 May of four consecutive years (2008-2011) on *G. illyricus* plants growing in the Sierra de Cazorla mountains (Jaén province, southeastern Spain). Every study year inflorescences were tagged before opening the first flower, and randomly assigned to either experimental or control groups. Experimental inflorescences were bagged with fine fabric mesh to preclude pollinator access and nectar colonization by yeasts, while allowing air to freely pass through and thus minimizing the build-up of internal excess humidity relative to the air outside. Control inflorescences were left exposed to natural visitation by pollinators, which could bring yeast inocula to floral nectar with their mouthparts. Tagged inflorescences remained in the field until each had at least its first 2-4 flowers open, at which time they were cut and brought to the laboratory in a portable cooler at 4-6°C. All open flowers in each inflorescence were labelled individually and their nodal position recorded (Figure 1A). Inflorescences or bags showing any sign of damage by deer or wild boar (mostly bagged ones) were discarded. Basal flowers with withering petals borne by a few inflorescences were also excluded. The final sample consisted of 219 flowers in 85 control inflorescences, and 153 flowers in 55 experimental inflorescences. The mean (± SE) nodal position of both the earliest (2.6 ± 0.1 and 2.9 ± 0.2) and latest (1.0 ± 0 and 1.1 ± 0.05) flowers were closely similar for control and experimental inflorescences, respectively, which denotes that flowers in the two treatment levels were of roughly similar age.

Nectar was extracted from each flower with calibrated microcapillaries and split into two subsamples. Total sugar concentration to the nearest 0.5% was determined in one subsample using a temperature-compensated, handheld refractometer (Bellingham & Stanley Ltd., Tunbridge Wells, UK). Nectar sugar concentration measurements were expressed as percent sucrose equivalents on a weight-to-weight basis. The other subsample was used for estimation of yeast cell density and assessment of yeast species composition. Nectar volume was calculated from the length of the nectar column within 5 µL or 10 µL calibrated pipettes (Drummond Scientific Company, Broomall, Pennsylvania) and it was then diluted up to 5–8 µL by addition of lactophenol cotton blue solution to facilitate microscopic examination. Yeast cell density in nectar (cells/mm^3^) was estimated under a microscope using a Neubauer chamber and standard cell counting methods.

For the purpose of yeast species identification, 20 nectar samples were collected in 2008 with sterile microcapillaries from as many untagged *G. illyricus* inflorescences from the study area which had been exposed to natural pollinator visitation. A 1-μL aliquot of each sample was streaked onto a Yeast Mold (YM) + chloramphenicol agar plate and incubated at 25°C. For each cultured nectar sample, distinct yeast isolates were obtained from all the resulting colonies following standard methods and criteria described in Yarrow (1998). For each isolate ∼500 nucleotides of the D1/D2 domain of the 26S subunit ribosomal DNA, the gene most commonly used for yeast identification, were two-way sequenced following methods in Kurtzman and Robnett (1998) and Lachance et al. (1999). The nucleotide collection databases at Genbank were queried with the Basic Local Alignment Search Tool (BLAST; Altschul et al. 1997) to look for described yeast species with DNA sequences matching those of isolates from *G. illyricus* nectar. *Metschnikowia reukaufii* and *M. gruessii* were the only yeast species identified (author, unpublished data). Free-living cells and colonies of these two species are morphologically distinct (Giménez-Jurado 1992; Kurtzman and Fell 1998) and can be easily distinguished microscopically, particularly on account of the unique “airplane or trident configuration” (Kurtzman and Fell 1998) shown by *M. gruessii* (Figure 1B and 1C). In addition to shape differences of cells and colonies, the two species stain differently with lactophenol: solid blue in *M. reukaufii*, only cell ends in *M. gruessii* (Figure 1B and 1C). All these differences allowed a reliable visual assessment of their relative abundances in the nectar samples examined microscopically. Each individual flower with yeasts was classed into one of the following four categories: (i) only *M. reukaufii*; (ii) both species present, *M. reukaufii* numerically dominant; (iii) both species present, *M. gruessii* numerically dominant; and (iv) only *M. gruessii*. In addition to yeasts, bacteria were isolated from *G*. *illyricus* nectar samples collected in lowland Mediterranean habitats ∼370 km away from the Cazorla study site (Álvarez-Pérez and Herrera 2013), but I found no microscopical evidence of bacteria in the nectar samples considered here.

The effect of experimental treatment (inflorescence bagging) on the within-inflorescence gradient in nectar sugar concentration was tested by fitting a linear mixed-effects model to data with nodal position, treatment and their interaction as fixed effects, and sampling year and inflorescence nested within year as random effects. Under this model design, a statistically significant interaction effect would denote that the within-inflorescence gradient in nectar concentration differed among treatment levels (control and bagged). The possibility that the two yeast species modified the vertical gradient of nectar concentration in different ways was examined by fitting a linear mixed-effects model to the subset of control flowers which had yeasts in nectar. In this model, nodal position, yeast species composition (as assessed with the relative abundance scale mentioned above) and their interaction were the fixed effects, and sampling year and inflorescence nested within year the random effects. A significant interaction term would reveal that variation in yeast species composition led to contrasting vertical gradients in sugar concentration within inflorescences. All statistical analyses were carried out using the R environment (version 4.6.0; R Core Team 2026).

Models were fitted using the lmer function in the package lme4 (Bates et al. 2015). Models were checked for linearity, homogeneity of variance and normality of residuals with function check_model in the performance package (Lüdecke et al. 2021), and goodness of fit was assessed with marginal (fixed effects only) and conditional (both fixed and random effects) *R*^2^ values estimated using function r2_nakagawa in the same package. The statistical significance of fixed effects in models was tested using Wald Chi-square tests as implemented in the Anova function of the car package (Fox and Weisberg 2019). Marginal effects and model-adjusted predictions of the response variable (sugar concentration) were estimated with the predict_response function of the ggeffects package (Lüdecke 2018).

## Results

In the set of flowers of control (exposed) inflorescences from all study years and nodal positions pooled, statistically significant trends emerged of increasing nectar sugar concentration (Chi-squared = 32.0, df = 3, *P* = 5.2E-07, Kruskal-Wallis rank sum test), and declining frequency of occurrence (Chi-squared = 21.5, df = 3, *P* = 8.1E-05) and density of yeasts (Chi-squared = 20.3, df = 3, *P* = 15.0E-05), from low to high nodal positions in inflorescences (Table 1). Bottom flowers (the oldest ones in each inflorescence) stood out from the rest by the high frequency of occurrence (80%) and density of yeasts in nectar (6,400 cells/mm^3^) (Table 1).

**Table 1.** Nectar sugar concentration, yeast frequency of occurrence and yeast cell density in field-sampled flowers of *Gladiolus illyricus* exposed to natural visitation by pollinators.

| Nodal<br>position in<br>inflorescence | Sample size<br>(number of<br>flowers) | Nectar sugar<br>concentration ±<br>SE (%) | Proportion of<br>flowers with<br>nectar yeasts ± SE | Yeast density (cells/mm<sup>3</sup>)<sup>a</sup> | |
| --- | --- | --- | --- | --- | --- |
| | | | | Mean ± SE | Interquartile<br>range |
| 1 | 85 | 22.3 ± 0.9 | 0.795 ± 0.04 | 6412 ± 1069 | 726-7493 |
| 2 | 76 | 26.1 ± 0.8 | 0.587 ± 0.06 | 3062 ± 577 | 486-4824 |
| 3 | 46 | 28.5 ± 0.9 | 0.415 ± 0.08 | 652 ± 162 | 144-967 |
| 4 | 12 | 27.1 ± 1.9 | 0.364 ± 0.15 | 227 ± 102 | 116-272 |

<sup>a</sup> : Computed only for the subset of sampled flowers whose nectar contained yeasts ( $N = 131$ ).

Not a single yeast cell was detected in nectar from flowers inside experimental (bagged) inflorescences. This corroborated expectations that pollinator visitation was responsible for yeast colonization of the floral nectar of *G. illyricus*, and that pollinator exclusion was effective at producing yeast-free experimental flowers that would exhibit plant-intrinsic gradients of nectar sugar concentration within inflorescences. The mixed-effects model testing the effects of nodal position, experimental treatment and their interaction on nectar sugar concentration fitted well the data, as shown by conditional (0.631, 95% confidence interval = 0.457–0.737) and marginal *R*^2^ (0.442, 0.310–0.540). There were highly significant effects of nodal position (Chi-squared = 19.1, df = 1, *P* = 1.2E-05), treatment (Chi-squared = 148.6, df = 1, *P* < 2.2E-16) and their interaction (Chi-squared = 83.8, df = 1, *P* < 2.2E-16) on nectar sugar concentration. Due to the significant node x treatment interaction each main effect is only interpretable in relation to the other (Figure 2A). Sugar concentration tended to decline from low to high nodal positions in the experimental inflorescences (yeasts excluded), while the reverse was true for control inflorescences exposed to natural pollinators (mostly with yeasts, Table 1). The strong interaction between nodal position and levels of the experimental treatment revealed an almost perfect mirror-image, symmetrical reversal by yeasts of the plant intrinsic vertical gradient, i.e., the one created by plants in absence of pollinators and associated yeasts (Figure 2A).

**Figure 2.**
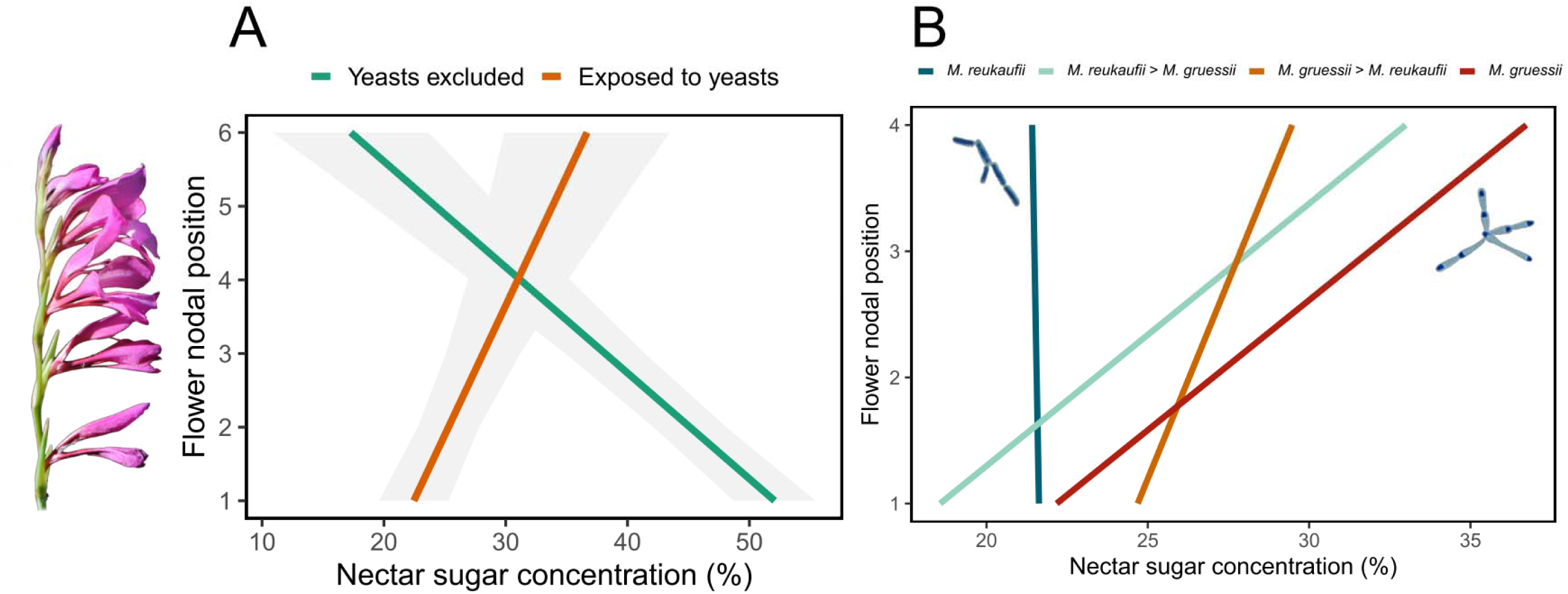
Interaction plots of the marginal effect of flower nodal position along *Gladiolus illyricus* inflorescences on nectar sugar concentration for inflorescences exposed to natural pollination and with pollinators excluded experimentally. **A.** Lines are predicted marginal effects of nodal position after accounting for variation among sampling years and inflorescences (included as random effects in model), shaded bands are 95% confidence intervals of predictions. **B.** Results for the subset of inflorescences exposed to natural pollination which had yeasts in nectar, separately for the four classes of yeast species composition. Lines are predicted marginal effects of nodal position after accounting for variation among years and inflorescences (included as random effects in model). Different ranges of nodal positions in **A** (1-6, flowers from all inflorescences) and **B** (1-4, only flowers with yeasts from exposed inflorescences) are due to a few bagged inflorescences bearing 1-2 long-lasting, old flowers at bottom.

The difference between *M. reukaufii* and *M. gruessii* in their impact on the vertical gradient in nectar sugar concentration was tested for the subset of *N* = 131 yeast-containing flowers. The linear mixed-effects model testing for effects of nodal position, yeast composition and their interaction on nectar sugar concentration fitted well the data (conditional and marginal *R*^2^ = 0.426 and 0.185, 95 % confidence intervals = 0.171–0.720 and 0.044–0.436, respectively). The effects of nodal position (Chi-squared = 9.8, df = 1, *P* = 0.0017), yeast species composition (Chi-squared = 8.6, df = 3, *P* = 0.035), and their interaction (Chi-squared = 9.5, df = 3, *P* = 0.024) were all statistically significant. The interaction plot in Figure 2B shows that *M. reukaufii*-only and *M. gruessii*-only classes had the weakest and strongest impacts on the slope of the vertical gradient of sugar concentration, respectively, and that mixtures of the two species had intermediate effects.

The impact of the nectar yeast community on the vertical sugar concentration gradient increased in the direction *M. reukafii* alone, *M. reukafii* dominant, *M. gruessii* dominant, and *M. gruessii* alone (Figure 2B).

## Discussion

When pollinators and associated yeasts were artificially excluded from the inflorescences of *G. illyricus*, a declining gradient in nectar sugar concentration built up which ran from the oldest, female-stage flowers at bottom up to the youngest, male-stage ones at top. The directionality of this gradient represents an intrinsic condition of plants arising without intervention of other organisms. This intrinsic gradient was fully reversed in inflorescences exposed to natural visitation by pollinators, in which sugar concentration increased from female-stage, oldest bottom flowers up to male-stage, youngest top flowers. This reversal entailed a shift in nectar sugar rewards between the sexual stages of protandrous *G. illyricus* flowers. Over the life of individual flowers, therefore, the highest energetic reward per nectar volume unit occurred during the late female stage (bottom positions) in unvisited experimental inflorescences, but during the early male stage (top positions) in inflorescences exposed to natural pollinator visitation and yeast colonization.

From a purely statistical perspective, the experimental design adopted in this study was unable to dissect the effects of pollinators and yeasts on nectar sugar concentration, since flowers exposed to pollinators were also exposed to yeasts by design. This confounding of effects was a irreducible biological reality of the study system, since the two species of *Metschnikowia* reported here are narrow nectar specialists invariably associated with large bees, particularly bumble bees (*Bombus*), and they colonize flower nectar when the latter is probed by pollinators with their mouthparts (Brysch-Herzberg 2004; Canto et al. 2008; Guzmán et al. 2013; Rutkowski et al. 2023). Regardless of strictly statistical considerations, however, some findings indicate that the experimental results of this study actually denote the action of yeasts. As a consequence of their intense sugar metabolism, the proliferation of *Metschnikowia* yeasts in nectar reduces total sugar concentration and sucrose-to-hexoses ratio of nectar, and originates inverse relationships of sugar concentration with flower age and yeast cell density (Canto et al. 2008; Herrera and Alonso 2025; Herrera et al. 2008; Schaeffer et al. 2015). For example, inoculation of artificial sucrose-dominated nectar with bumblebee-vectored *Metschnikowia* yeasts almost depleted sucrose and reduced sucrose-to-hexoses ratio from 51-67 to 2-6 in 48 h (Canto et al. 2008). The inverse relationships found here along nodal positions of exposed flowers between nectar sugar concentration, on one side, and flower age and yeast cell density on the other, therefore confirm expectations from the sugar-degrading metabolism of nectar yeasts (Herrera et al. 2008). The species-specific effect on inflorescence-wide sugar concentration gradients likewise supports the causal role of yeasts, since *M. reukaufii* and *M. gruessii* differ widely in physiological profiles (Giménez-Jurado 1992; Kurtzman and Fell 1998; Pozo et al. 2016). Of particular relevance here is the fact that *M. gruessii* cells multiplicate much faster than those of *M. reukaufii* in sucrose-dominated substrates such as *G. illyricus* nectar (Pozo et al. 2016).

Earlier studies on functional correlates of the Darwin’s inflorescence syndrome (see Introduction for background and references) have focused infrequently on vertical gradients of nectar sugar concentration, and previous investigations do not seem to have compared within-inflorescence patterns of nectar sugar concentration between inflorescences with and without pollinators as done here. The mixed results of the few investigations available, however, make sense in the light of findings presented here. In five *Digitalis* species studied by Percival and Morgan (1965), nectar concentration in flowers (apparently) exposed to visitation by pollinators consistently increased from base to top of inflorescences, as found here for openly pollinated flowers. In contrast, variation in nectar concentration within bagged inflorescences was either nonsignificant (Pyke 1978; Roguz et al. 2025) or declined from bottom to top (Best and Bierzychudek 1982; Galen and Plowright 1985), as found here for bagged *G. illyricus* inflorescences. This disagreement among previous studies could reflect their heterogeneity with regard to the presence of pollinator-borne yeasts in the flowers studied. *Metschnikowia* yeasts, in addition to being widespread worldwide in nectar of entomophilous flowers, are mainly associated with bees (Brysch-Herzberg 2004; Canto and Herrera 2012; Canto et al. 2017; de Vega et al. 2009; Herrera et al. 2008, 2009; Lachance et al. 2001; Rutkowski et al. 2023), the main pollinators of plants with Darwin’s inflorescence syndrome (Strelin et al. 2024). I tentatively suggest that yeast inflorescence engineering similar to that experimentally documented in this note for *G. illyricus* might eventually prove to be widespread among bee-pollinated plants with Darwin’s inflorescence syndrome whose flowers are colonized by nectar yeasts, particularly by those with faster multiplication rates. Given that the relative multiplication rates of *M. reukaufii* and *M. gruessii* in nectar depend on the latter’s sucrose-to-hexoses ratio (Pozo et al. 2015), the inflorescence engineering ability of yeasts should be expected to depend on the combination of dominant yeast species and plant-specific nectar sugar composition.

The presence of *Metschnikowia* yeasts in nectar of individual flowers enhances floral signalling, flower attractiveness and visitation rate by bee pollinators, a phenomenon presumably mediated by modification of floral scent due to volatile emission by yeasts (Deng et al. 2024; Herrera et al. 2013; Rering et al. 2018; Schaeffer et al. 2014, 2017, 2019; Yang et al. 2019). For instance, bumblebee visitation to flowers of *Clematis akebioides* was higher to flowers with *M. reukaufii* yeasts, which emit 2-methyl-1-butanol, an electro-antenographically compound active for bees (Rering et al. 2017; Yang et al. 2019). As these studies have consistently documented a foraging preference of bees for yeast-containing flowers, the high densities attained by *Metschnikowia* in the oldest, lowermost flowers of *G. illyricus* inflorescences exposed to pollinators should represent an attractive, self-advertising target when bees first approach an inflorescence. Through associative learning (Schaeffer et al. 2017), such hypothesized effect should reinforce the starting preference of bees for bottom, yeast-rich flowers, to be followed next by the upward displacement which characterizes bee foraging on Darwin’s syndrome inflorescences. The vertical displacement from low reward but attractive bottom flowers to the progressively less attractive but more rewarding ones at upper nodal positions should be favored by the positive responses of bees to nectar sugar concentration (Cnaani et al. 2006; Whitney et al. 2008). In this way, and despite the microbial reversal of the vertical reward gradient, yeast-engineered *G. illyricus* inflorescences would still fit the "female first" vertical foraging directionality which is central to the Darwin’s inflorescence syndrome.

A rigorous assessment of the impact of yeasts on pollination success and pollen carryover within and among inflorescences (de Vega et al. 2022) was beyond the scope of this study, as it would require experiments comparing populations with and without floral yeasts.

Populations of *G. illyricus* without yeasts do not seem to exist in my study area, as *Metschnikowia* yeasts were found in flowers of the *N* = 15 populations examined in a pilot survey conducted there in 2008 (author, unpublished data). Even in absence of information on reproductive consequences for plants, however, yeast reversal of intrinsic nectar sugar gradients within Darwin’s syndrome inflorescences reported here can have implications in relation to the inferred ecological and evolutionary significance of inflorescence organization (Harder et al. 2004). In particular, it can be predicted that the differences found here between *M. reukaufii* and *M. gruessii* in their effects on the sugar concentration–nodal position slope should result in measurable differences in within-inflorescence pollen flow and, accordingly, variation in progeny outcrossing rate. The variable inflorescence engineering ability of cosmopolitan *Metschnikowia* yeasts could account for variable results in experiments of pollinator foraging and inflorescence functionality depending on abundance, species composition and within-inflorescence distribution of yeasts in the flowers. As floricolous *Metschnikowia* yeasts are more closely associated with bees than with other pollinators, and bees are in turn associated with the Darwin’s inflorescence syndrome, the possibility exists that yeasts could be playing an hitherto unrecognized role in shaping the vertical profiles of nectar sugar chemistry and concentration along this type of linear inflorescences. More generally, results presented in this note support the notion that inflorescence traits typically attributed to plant genotype and pollinator-mediated selection can actually reflect the metabolism of ubiquitous nectar yeasts. When these microbial effects go unmeasured in observational or experimental field studies, phenotypic patterns can be misattributed to pollinator selection rather than a direct consequence of microbial presence.

## Ethics

This work did not require ethical approval from a human subject or animal welfare committee.

## Data accessibility

Data, metadata and R script used for analyses are available for editors and reviewers at figshare, https://figshare.com/s/25909931e52ccd930d47

## Declaration of AI use

I have not used AI-assisted technologies in creating this article.

## Acknowledgements

I am indebted to Pilar Bazaga and Esmeralda López-Perea for assistance with molecular identification of yeasts; Judith Bronstein and two anonymous reviewers for comments and suggestions that improved considerably this note; and Consejería de Medio Ambiente, Junta de Andalucía por permission to work in the Sierra de Cazorla and providing invaluable facilities there.

